# Sustained EEG responses to rapidly unfolding stochastic sounds reflect precision tracking

**DOI:** 10.1101/2024.01.08.574691

**Authors:** Sijia Zhao, Benjamin Skerritt-Davis, Mounya Elhilali, Frederic Dick, Maria Chait

## Abstract

The brain is increasingly viewed as a statistical learning machine, where our sensations and decisions arise from the intricate interplay between bottom-up sensory signals and constantly changing expectations regarding the surrounding world. Which statistics does the brain track while monitoring the rapid progression of sensory information?

Here, by combining EEG (three experiments N≥22 each) and computational modelling, we examined how the brain processes rapid and stochastic sound sequences that simulate key aspects of dynamic sensory environments. Passively listening participants were exposed to structured tone-pip arrangements that contained transitions between a range of stochastic patterns. Predictions were guided by a Bayesian predictive inference model. We demonstrate that listeners automatically track the statistics of unfolding sounds, even when these are irrelevant to behaviour. Transitions between sequence patterns drove an increase of the sustained EEG response. This was observed to a range of distributional statistics, and even in situations where behavioural detection of these transitions was at floor. These observations suggest that the modulation of the EEG sustained response reflects a universal process of belief updating within the brain. By establishing a connection between the outputs of the computational model and the observed brain responses, we demonstrate that the dynamics of these transition-related responses align with the tracking of ‘precision’ – the confidence or reliability assigned to a predicted sensory signal - shedding light on the intricate interplay between the brain’s statistical tracking mechanisms and its response dynamics.

## Introduction

The brain is increasingly conceptualized and modelled as a statistical learning machine. On this account, our sensations and decisions arise via an interaction of bottom-up sensory signals with continuously updated expectations (“model beliefs”) about our surrounding environment. A major current research effort in systems neuroscience is to uncover the properties of this “observe-predict-update” loop. Fundamental to this goal is understanding how the statistics of the environment are being monitored and represented by perceptual systems.

Based on Bayesian ideal observer models, theoretical accounts of these inference processes suggest that in noisy and volatile environments, optimal belief updating depends on the observer’s ability to assess both the stability and the reliability of the environment (1–4). Specifically, observers are hypothesized to track the probability that the environment has changed, i.e., that the generative process underlying the observations is now different (1,5–7). They must also simultaneously update a predictive model of the next observation, based on previously encountered inputs. Evidence suggests that the latter process involves not only the representation and continuous updating of the next expected input, but also tracking of the confidence, or *inferred reliability* (“precision”), of this prediction., such that incoming events are evaluated based on this weighing (8–14).

Belief updating has predominantly been studied in research on probabilistic decision making. Using relatively slow tasks that depend on evidence accumulation, multiple studies have shown that participant responses reflect estimated precision (e.g., 2,15,16). For example, in a seminal experiment (2) participants performed a task that required them to predict the next number in a series. The numbers were drawn from a Gaussian distribution with a mean that changed abruptly at random intervals. Results indicated that performance and baseline pupil diameter - hypothesized to index arousal and autonomic state - were modulated by belief uncertainty, as modelled with a Bayesian model of evidence accumulation. More recently, an EEG study (13) showed that the cortical activity tracks word surprisal in continuous natural speech, and that this tracking is modulated by precision. Here we focus on understanding evidence accumulation in a faster-paced domain, asking what statistics of rapidly unfolding sensory signals are automatically tracked by the brain during listening.

The acoustic environment is rich in statistical regularities that unfold on many different time scales. These statistics characterize auditory textures (water, fire, a roaring engine, the hum of a crowd), reflect physical constraints of sources (locomotion sounds, vocalization), and register the reoccurrence of events across time (the repeating call of a bird in the forest; or motifs in music). The auditory system continuously analyses and tracks these regularities as they appear and disappear from our surroundings - even when this information is not immediately relevant to behavior (6,17–20). Experimentally manipulating the statistical structure of auditory sequences therefore opens important windows onto the brain’s statistical learning machinery. In particular, it offers a means to probe potential heuristics used by the system, to ascertain which statistical information is tracked, and to disentangle more ‘automatic’ processes from those related to carrying out behavioural goals.

In this vein, observations from paradigms based on introducing deviants in structured sound sequences have shown that deviant-evoked brain responses reflect precision-weighted prediction errors (10,21–25). Recent dynamic causal modelling (DCM) work has suggested that encoding of precision and prediction errors may be dissociated (26).

Nevertheless, deviant-evoked responses provide only an indirect and momentary measure of listeners’ sensitivity to ongoing statistical structure. To circumvent this issue, Barascud et al. (27) used the dynamics of brain responses to statistically structured auditory streams to investigate the process by which listeners automatically acquire an internal model of regularities in the environment. Here, naïve distracted listeners were presented with sequences of short tone-pips containing transitions from randomly ordered to regularly repeating patterns. The emergence of regularity was associated with a prototypical brain response pattern: a gradual increase followed by a plateau in sustained EEG power, one generated by auditory cortical, frontal, and hippocampal sources. Increases in sustained EEG power toggled to increases in sequence predictability have been observed for a variety of patterned stimuli (17,28–31). The greater EEG amplitude for regular over random stimuli cannot easily be interpreted as a perceptual response to physical attributes of the signal. Adaptation, for example, would be expected to result in the opposite pattern (32,33). Rather, Barascud et al. (27) showed that the sustained response appears to vary consistently with the predictability of the ongoing stimulus sequence and hypothesized that it might reflect inferred reliability (“precision”) (34).

However, this previous work suffers from a major limitation: these studies only used transitions from random to fully deterministic sequences, i.e., precisely repeating patterns. This makes it difficult to disambiguate the predictions made under a Bayesian framework - emphasizing the process of evidence accumulation under uncertainty, and distinguishing “change probability” tracking from “precision tracking” - from those from memory-based accounts, which emphasize mechanisms engaged by the repeating tone sequences (7,35,36).

Here, we combined EEG and computational modelling to investigate brain tracking of rapid, stochastic sound sequences. Passively listening participants were exposed to structured tone-pip arrangements (Figure 1) that contained transitions between a range of stochastic patterns, manifested as abrupt changes in the mean and/or variance of the unfolding sequence. Importantly, the sequences did not contain transient deviants; distributional changes can only be inferred by tracking the unfolding sequence statistics. This thus allows us to ask directed questions concerning what statistical information about the sequence structure is reflected in EEG response dynamics.

**Figure 1.**
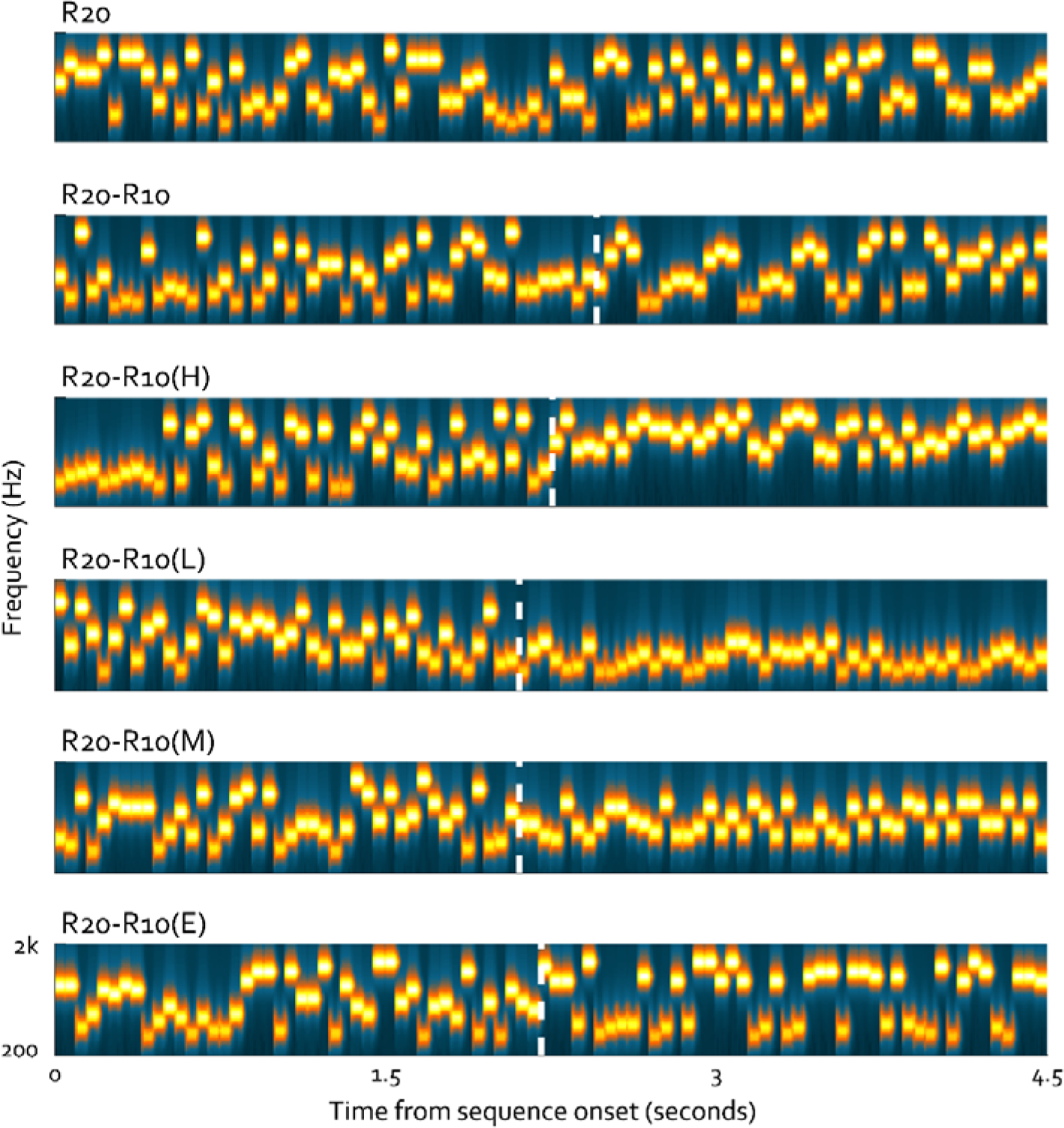
Example spectrograms of stimuli in Experiment 1. The stimuli were sequences of concatenated tone-pips (50ms) with frequencies drawn from a pool of 20 fixed values, spaced from 222 Hz to 2000 Hz in steps of ∼2 semitones. Six different conditions were used. The control condition, R20, was generated by randomly sampling from the full pool with replacement; The remaining conditions contained a change partway through the trial, manifested as a reduction in alphabet size from 20 to 10 tones. In R20-R10 the retained tones were sampled equally from the full pool. In R20-R10(H) the retained tones were sampled from the two highest frequency bands (the 10 highest frequencies; 707Hz – 2000Hz). In R20-R10(L) the retained tones were sampled from the two lowest frequency bands (10 lowest frequencies; 222Hz – 630 Hz). In R20-R10(M) the retained tones were sampled from the two central bands (10 middle frequencies; 397Hz – 1122 Hz). In R20-R10(E) the retained tones were sampled from the highest (1259-2000Hz) and lowest (222-354 Hz) bands (5 highest and 5 lowest frequencies). For illustrative purposes, the plotted sequences are of a fixed length. In the actual experiment, transitions (marked as dashed white lines here) occurred between 2 to 2.5 seconds post sequence onset, with the overall sequence length varying from 4 to 4.5 seconds.

Predictions were guided by a Bayesian predictive inference model (37; Figure 2) previously used to study a variety of statistical tracking phenomena (1,2,16). To explain the behaviour of human observers in volatile environments, the model monitors sequences containing hidden changes in the underlying statistical structure. As it “listens” to the unfolding auditory sequence, it derives a predictive distribution for the next observation using sufficient statistics collected over a local context of previous observations, the extent of which adapts to the volatility of the observed sequence. The model monitors two key inferred statistics to guide predictions about upcoming input: (1) *change probability,* the likelihood that a change in sequence statistics has occurred; and (2) *precision* (inferred reliability), tracking how variable inputs have been in the relevant past. As highlighted above, sensitivity to both change probability and precision is hypothesized to play a fundamental role in perception. Accurate estimation of change probability is critical for survival in natural environments that often change abruptly and unpredictably. Tracking of precision is hypothesized to be critical to observers’ ability to operate under different degrees of uncertainty (8,38–40).

**Figure 2.**
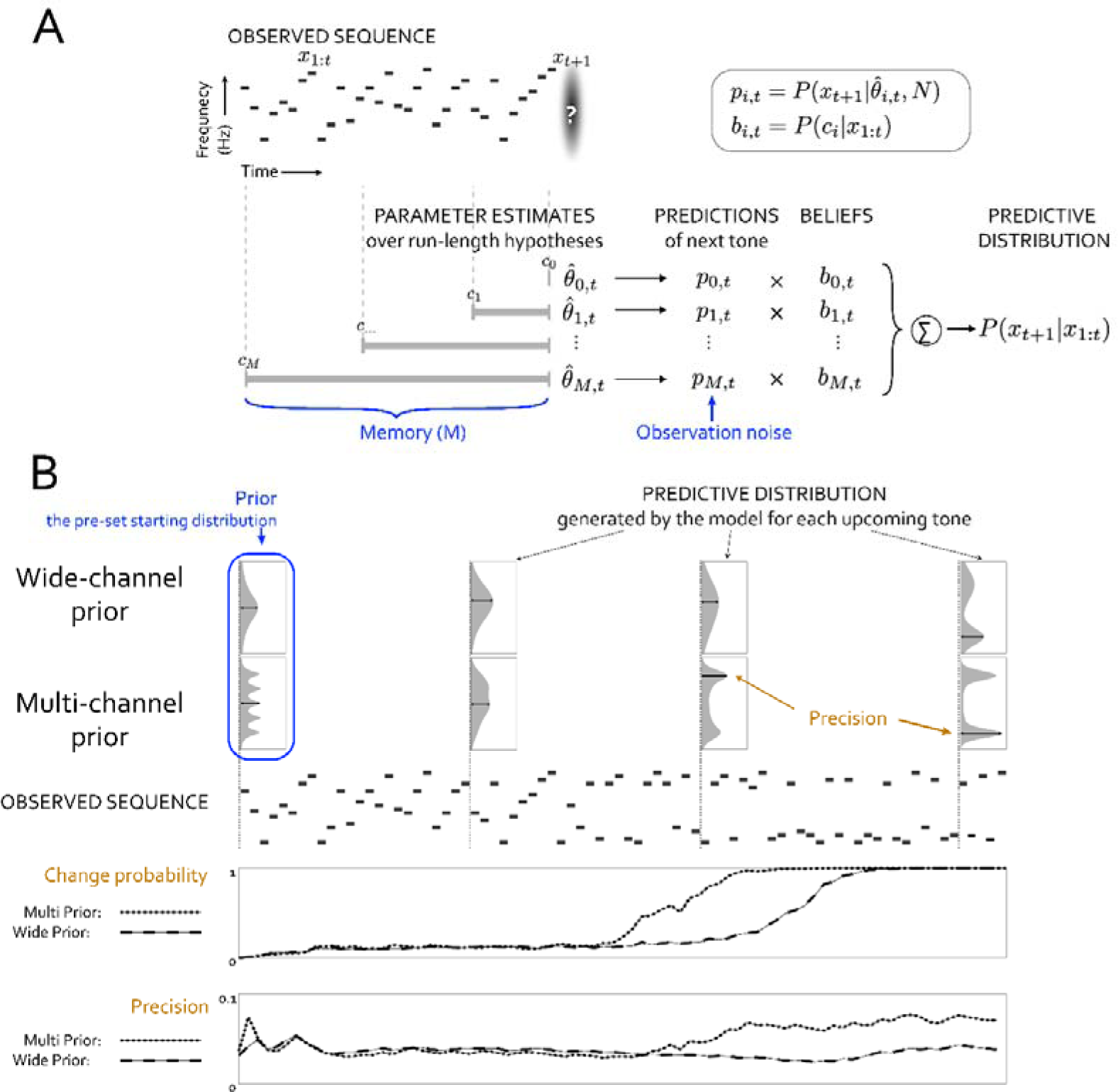
Schematic of perceptual model and model outputs. (A) At time *t*, the model contains multiple parameter estimates collected over run-lengths into the past up to the memory constraint (*M*). Each estimate yields a prediction for the next tone, with increased uncertainty due to observation noise. Upon observing *x_t+1_*, the model updates the run-length beliefs using the predictive probability for each hypothesis. Note that the prediction for length *M* is used to update all beliefs with length greater than or equal to *M*, thus limiting the number of past observations used in the update. (B) Outputs from the model for an example observed sequence (here the unfolding stimulus is R20-R10(E)). The predictive distribution (Gaussian distribution shown on the top), generated by the model for each upcoming tone, combines predictions across run-length hypotheses weighted by their beliefs, thus “integrating out” run-length. At the beginning of each trial, a starting distribution (‘*Prior’)*, is initiated; Two prior types were modelled - a single Gaussian across the entire spectral range (wide-channel prior) or a Gaussian with multiple components (multi-channel prior). The change probability is the probability that at least one changepoint has occurred, as inferred using the run-length beliefs. To adapt the model to behavioural responses, the model detects a change if the change probability exceeds threshold. The precision is the maximum value of the normalised predictive distribution (marked as a black line in each predictive distribution). In this example, the change in stimulus statistics is associated with a dynamic increase in change probability and precision, which occurs earlier in the multi-channel prior model. The model parameters (memory, observation noise and priors) are in blue. The outputs (change probability and precision) are plotted in the bottom for each prior distribution (wide- and multi-channel prior).

By relating model outputs to observed brain responses, we asked (a) whether EEG responses exhibit sensitivity to the evolving structural patterns within auditory sequences; and (b) whether the dynamics of these transition-related responses align with change point estimation, or alternatively tracking of precision (as hypothesized in Barascud et al. (27)).

## Results

### Listeners are sensitive to the statistics of stochastic tone patterns

We developed a new stimulus paradigm, based on the tone patterns used in Barascud et al. (27), to study sequences that contain transitions between stochastic patterns that are differentially statistically structured (i.e., sequences that contain abrupt changes to the underlying generative parameters). Our basic stimulus, a random pattern of 20 frequencies (R20; an example is shown in Figure 1), consists of a rapid sequence of 50ms tone-pips whose frequencies are uniformly sampled from a discrete pool of 20 values (see Methods). We created transitions from the full pool of 20 values to another random sequence, but where the tones are drawn from a reduced set (“alphabet size”) of 10 frequencies (R20-R10). To address the core question - whether listeners are sensitive to changes in distributional statistics - we manipulated R20-R10 such that the 10 retained frequencies are sampled from different distributions (Figure 1).

The full pool of 20 frequencies was divided into four logarithmically equally sized frequency bands, each containing five frequency values. All transitions featured a reduction to 10 frequencies. These were selected from (a) the full pool of frequencies (R20-R10); (b) the highest (R20-R10(H)) spectral band; (c) lowest band (R20-R10(L)); (d) middle band (R20-R10(M)); and (e) edge bands (R20-10(E); see the figure legend of Figure 1 and Methods for details).

**R20-R10(H)** and **R20-R10(L)** cause a change in both the mean and variance of the spectral distribution, and thus should evoke a transition to a single-peaked spectral predictive distribution. **R20-R10(M)** drives only a change in variance, but again a single-peaked spectral distribution. **R20-R10(E)** will drive only a change in variance, but a more complex, double-peaked spectral distribution. Finally, **R20-R10** is only associated with a reduction in alphabet size, but neither a change in mean nor variance.

As an initial step to characterizing listeners’ sensitivity to such changes in spectral statistics over time, we ran a behavioural study where participants were instructed to press a button immediately upon hearing a transition within a sound sequence. To allow participants to develop the most appropriate listening strategy, each stimulus type was presented in a separate block. In each block, 20 trials of the transition condition were randomly intermixed with 20 trials of its non-change control (R20). The order of the five blocks was randomised across participants.

In principle, there are at least two broad strategies which listeners could employ in the process of transition detection: (1) tracking the zeroth order statistics of the unfolding sequence (i.e., the probability of each frequency); (2) monitoring the distributional statistics of tone frequencies (i.e,. the probability distribution across the spectrum), either by tracking *a wide-band* (cross spectral) distribution covering the entire spectral range, or by tracking several concurrent (*’multiband’*) spectral distributions reflecting activity in distinct spectral channels. In (1) the transition will manifest as an increase in incidence in those channels (from probability of 1/20 to 1/10) that are retained after the transition, and in a decrease in incidence in the rest of the channels (from a probability of 1/20 to 0). If listeners do employ *Strategy 1*, we expect no difference in performance across conditions because the same number of tones (frequency channels) are retained in each. In contrast, *Strategy 2* would predict graded performance across conditions as a function of the similarity between pre/post transition means and variances. Here, R20-R10(H) and R20-R10(L) both contain changes in the average and variance and hence should be detected most readily, with somewhat poorer detection in R20-R10(M) (change in variance only). R20-R10(E) transitions should be yet more difficult to detect because they are associated with the emergence of a double peaked post-transition distribution. Finally, R20-R10 should be most difficult because there is no change in variance or mean.

Figure 3A summarizes the behavioural performance. A repeated measures ANOVA on d’ data showed a main effect of condition (*F*(4, 36) = 32.99, *p* < 0.001). Post-hoc tests revealed that the *d’* for R20-R10 was significantly lower than all other conditions (*p* ≤ 0.001). The second lowest condition was R20-R10(E) followed by R20-R10(M) (p = 0.022). The highest *d’* values were associated with R20-R10(H) and R20-R10(L), which did not statistically differ from each other (*p* = 0.60) but were significantly higher than for the other transition types (*p* ≤ 0.003). One-sample t-tests (2-tailed) for comparisons against 0 were applied to the *d’* data. Results established that performance on the H, L, M and E conditions was significantly above floor (p<0.001), whilst that for R20-R10 did not differ from floor (t(9) =0.76, p = 0.47).

**Figure 3.**
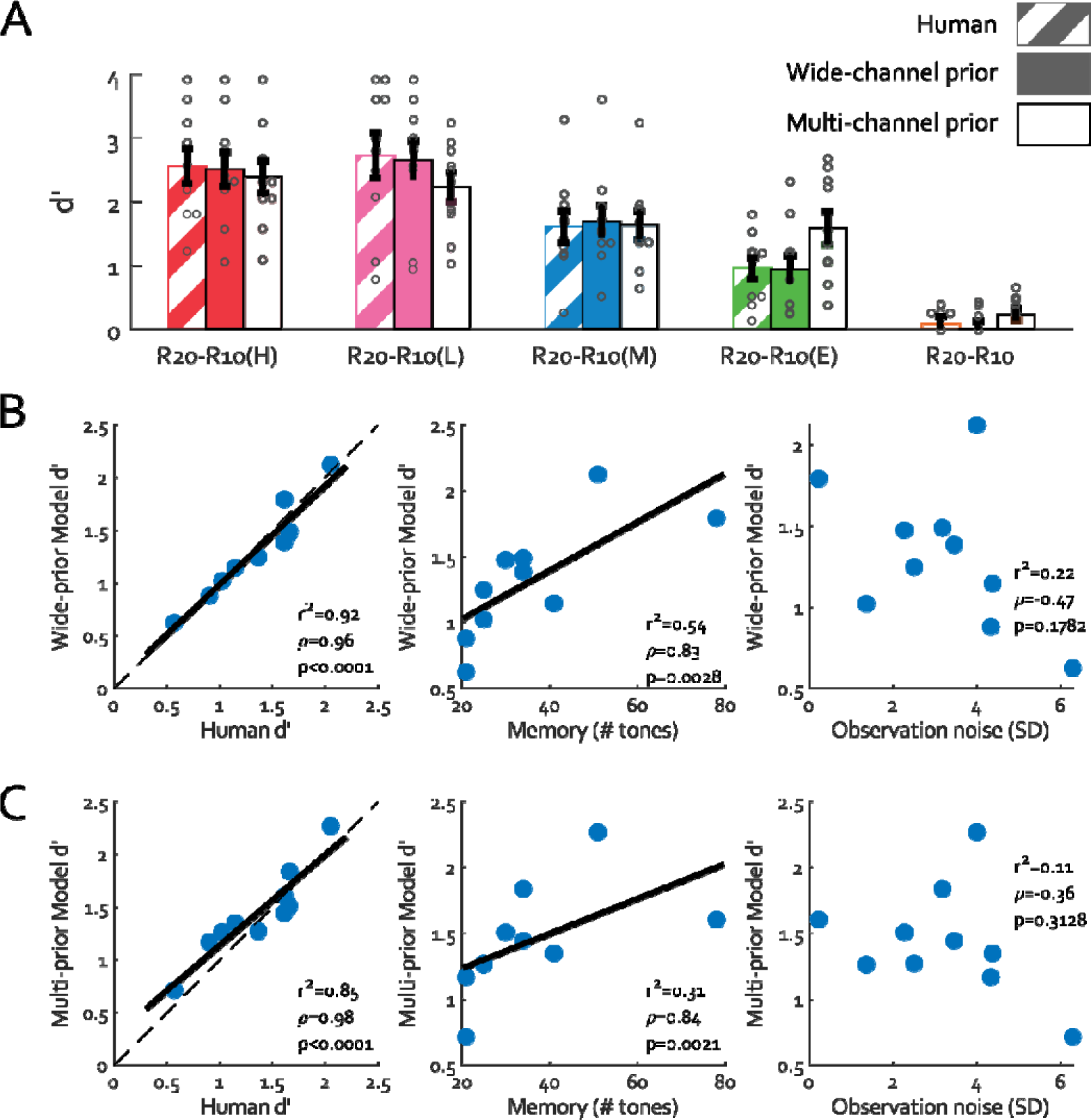
Behavioural and modelling results for Experiment 1, with results from humans (striped bars) and the D-REX models (solid bars, dark for wide-channel prior and light for multi-channel prior models). Coloured bars are group mean values, with grey circles indicating individual means. All error bars are 1 SEM. (B) & (C) Overall fit between human and model performance. Correlations between human and modelling results indicate that the D-REX model with wide-channel prior (B) and multi-channel prior (C) provides a good fit for human performance on an individual level. (Left) Human *d’* plotted against the model *d’*. Blue dots indicate individual participants. The black line indicates the linear fit of the individual data. Spearman’s correlation values are shown at the bottom right corner of the graph. Overall *d’* was computed by collapsing across all five transition conditions. We then examined the relationship between model performance (*d’*) and two parameters derived from the model fit to each participant: memory (M) and observation noise. (Middle) Like the relationship with the actual human *d’*, memory size is positively correlated with the model *d’*, while observation noise is not.

This results pattern demonstrates that behavioural performance is consistent with *Strategy* 2, where listeners track the distributional statistics of tone frequencies. Detection of R20-R10(H) and R20-R10(L) was most accurate, followed by R20-R10(M), 20-R10(E), with R20-R10 being least accurate.

### A Bayesian prediction model, D-REX, accurately fits behavioural data

We used a Bayesian sequential prediction model (Dynamic Regularity Extraction D-REX) (37) to fit the behavioural data (Figure 2).

As it is exposed to the unfolding tone-pip sequence, D-REX sequentially generates a predictive distribution (mixture of Gaussians) of the frequency of the upcoming tone given the previous observations (Figure 2A). At the beginning of each trial, the seed distribution is pre-set to represent the hypothetical listener’s default assumptions about the source distribution. Two classes of assumption were modelled (Figure 2B): 1), a *Wide-channel prior* modelled using a single Gaussian centred on the log-frequency range of the stimuli; or 2), a *Multi-channel prior* modelled as a multimodal Gaussian distribution. These priors represent two schemes for listening strategy #2, as discussed above: (a) listeners employ a *single wide-band* (cross spectral) distribution covering the entire spectral range; or (b) listeners employ a *within-channel* listening strategy, approximating a uniform distribution over frequency where several concurrent, partially overlapping distributions are being tracked simultaneously. To capture human listeners’ limited perceptual capacity, the model incorporates two perceptually plausible constraints: a finite memory, constraining the maximum number of previous observations used to generate the predictive distribution; and observation noise, representing individual differences in pitch perception.

We applied the model to the stimuli in Experiment 1 to fit human listeners’ behavioural performance. For each participant, the model was run on the entire experimental session in the same way they were delivered to the human participants. The comparison between human and model performance for each transition condition is shown in Figure 3A.

A repeated measures ANOVA on model *d’* data showed a main effect of condition for both the wide-prior model (*F*(2.82, 25.36) = 33.48, *p* < 0.001, η^2^= 0.79) and the multi-prior model (F(2.29,20.62) = 23.46, p < 0.001, η^2^ = 0.72). *Post-hoc* tests for the wide-prior model revealed that the *d’* for R20-R10 was significantly lower than all other conditions (p = 0.016 when compared with R20-R10€ and *p* < 0.001 compared with R20-R10(H)/(L)/(M)). Overall, the pattern was similar to that observed in humans: the second lowest *d’* was R20-R10(E), followed by R20-R10(M) (mean difference = 0.75 ± 0.27, p = 0.047). The highest *d’* values were associated with R20-R10(H) and R20-R10(L) which did not different statistically (*p* = 1.00) but were significantly higher than the other transition types (wide-prior: *p* ≤ 0.043). One-sample t-tests (2-tailed) for comparisons against 0 were applied to the *d’* data. Again, the wide-prior model was very similar to human performers in that it could not accurately detect the change in R20-R10 (not different from floor, t(9)=0.15, p = 0.89). Post-hoc tests for the multi-prior model revealed a slightly different pattern: unlike the single-prior model, a one sample t-test against 0 showed above chance performance for R20-R10 (t(9)=2.52, p = 0.033). Furthermore, the multi-prior model showed no difference between R20-R10(M) and R20-R10(E) conditions, outperforming human listeners in the latter.

Overall d’ was computed by collapsing across all five transition conditions. As shown in Figure 3B,C the model provides a good fit to (mean) human performance (wide-prior:, r^2^ = 0.92, p<0.0001; multi-prior: r^2^ = 0.85, p<0.0001). We further examined the relationship between human performance and two additional model perceptual parameters: memory and observation noise. The human performance (d’) is positively correlated with memory (wide-prior: r^2^=0.54, p = 0.028; multi-prior: r^2^=0.31, p = 0.021), suggesting that listeners modelled to have longer integration horizons were empirically better able to detect sequence transitions. No correlation with observation noise was observed (Figure3C right). Overall, this model, and in particular the ‘wide-channel’ prior, sensitized to unimodal distributional statistics, successfully captured human performance across a range of transition types. It also demonstrated that most listeners based their transition detection on a context length of 20-40 tones (1-2 seconds).

### The human brain automatically tracks the statistics of stochastic acoustic patterns

The initial data above are based on behavioural responses to perceived changes in sequence characteristics. To understand the more general case, we turned to EEG to ask whether the brains of ***naïve distracted listeners*** are sensitive to changes in stochastic acoustic patterns - and if so, what sequence statistics are being monitored. In Experiment 2A, participants listened to the R20-R10(H), R20-R10(M), R20-R10(E) transition conditions, along with R20 as a no-change control (50% of trials). (Unlike the behavioural task, all conditions were presented in random order, trial-by-trial). The R20-R10(L) condition was omitted to reduce experiment length, due to its similarity to R20-R10(H). In Experiment 2B, listeners listened to the R20-R10 (full spectrum) condition along with R20 as a control. In all experiment that follow, participants were naïve to the stimuli and watched a silent movie while listening.

Based on previous results showing increases in the sustained EEG power with increases in regularity (27–31), we expected that listeners’ sensitivity to statistical transitions would be reflected in an EEG divergence between the responses to R20-R10/H/M/E versus no-change R20.

As shown in Figure 4, all conditions evoked a similar EEG response: an N1-P2 onset complex followed by a persisting sustained response, and an offset response (see the topographies in Figure 4A). In the four change conditions, clear evoked responses are observed post-transition (Figure 4, A-D). The sustained response gradually diverged from R20 after the transition. Topographies (Figure 4F) show that this amplitude increase in sustained response is attributed to a steady rise of negative activity in central channels, consistent with effects observed previously (27,28). As discussed also elsewhere (27,28), the divergence between conditions is not outwardly consistent with low level adaptation effects. An adaptation-based explanation would in fact predict the opposite pattern: R10(H/M/E) sequences are associated with a *higher* probability of occurrence per tone, and hence should cause *greater* adaptation than R20.

**Figure 4.**
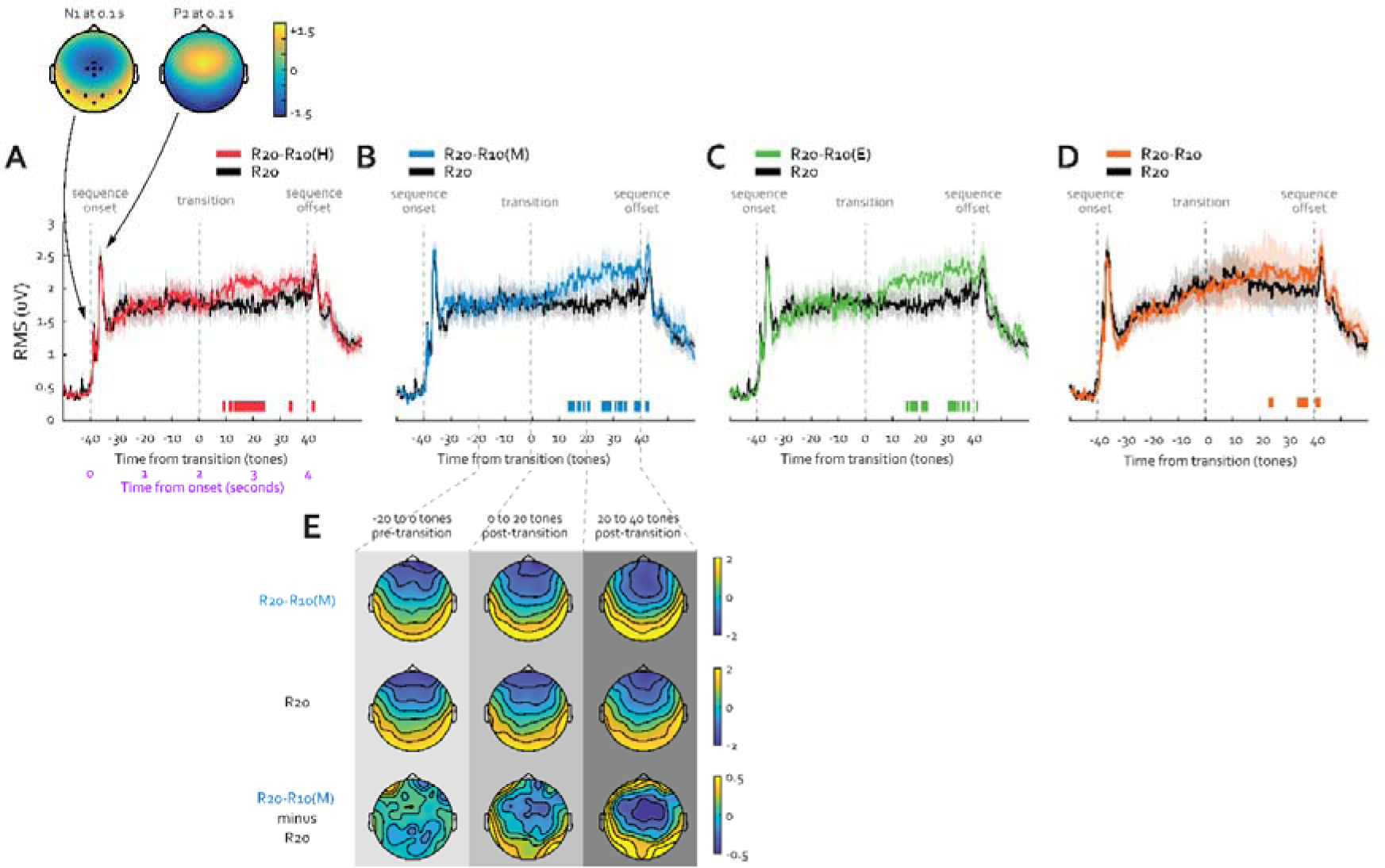
EEG results of Experiments 2A and 2B. (A-D) Plotted are the group mean of EEG responses from naïve distracted participants listening to sequences while watching a silent film. Root mean square (RMS) was computed for each participant over 10 auditory sensitive electrodes, selected based on the N1 response evoked by the sequence onset (left topography in A). Shaded areas are +/− 1 SEM (standard error of the mean). Colour-coded horizontal lines at graph bottom indicate time intervals where bootstrap statistics show significant differences between each transition condition and R20 (black trace). All trials had a fixed total length of 4 seconds (80 tones), with a potential (p = 0.5) transition in tone pool size starting at 2 seconds. The sequence onset, transition and offset are all marked with grey vertical dashed lines. Data in (A) to (C) are from Experiment 2A (N=26) and data in (D) is from Experiment 2B (a different group of participants; N=26). (E) Topographies for R20-R10(M) and the control R20 are plotted over three key time periods: (1) 1 second before transition (−20th to 0th tone pre-transition) (2) 1 second after transition (1st to 20th tone post-transition) and (3) the last second before offset (21st to 40th tone).

The sustained response amplitude did not differ across conditions (Repeated measures ANOVA on mean power post-transition between H, M and E; F(1.60, 39.98) = 0.17, p = 0.80, η^2^ = 0.007). Considering the behavioural detection performance as an indicator of the salience of the pattern change, the absence of variation in the sustained response amplitude among these three conditions implies that the sustained response is likely not a direct reflection of the behavioural salience of the transition itself. Instead, as will be shown in Experiment 3, below, it appears to be linked to the informational content inherent in the post-transition sequence.

### The EEG sustained response in passively listening, distracted individuals reflects sequence statistics

A potential explanation for the observed effects in Experiment 2 is that the amplitude of the sustained response is not associated with stimulus statistics *per se,* but instead reflects a heightened attentional state triggered by the *change* in the unfolding sequence. (Note that transitions were always task irrelevant, and participant attention was directed toward a silent movie). To examine this possibility, Experiment 3 selected one of the conditions – R20-R10(M), our “anchor condition” - and introduced a *reverse* transition R10(M)-R20, where the spectral distribution changes from reduced to full. (The stimulus set also included two no-transition conditions: R10M and R20).

If the gradual increase in the sustained EEG response merely reflects *detection of a statistical change,* then responses to R10M-R20 and R20-R10M should be similar because both contain statistical changes partway through the sequence. However, if the increased EEG power *is specific to the statistical properties of the spectral content before and after the transition*, R10M-R20 should evoke a different response compared to R20-R10M.

The latter predictions held (Figure 5). The response to R20-R10(M) shows a slow rise in the sustained response (Figure 5A), replicating the pattern observed in Experiment 2. The *opposite* pattern is observed in the response to the R10(M)-R20 transition, relative to its no-change control R10(M) (Figure 5B). Shortly after the R10(M)-R20 transition, there is an MMN-like response, corresponding to the presentation of a novel frequency (the first tone of the R20 portion of the sequence, see the high-pass filtered response at the bottom of Figure 5B). Following this response, a sharp *drop* in the sustained response is observed, later settling at a level below that of R10(M) for the remainder of the trial.

**Figure 5.**
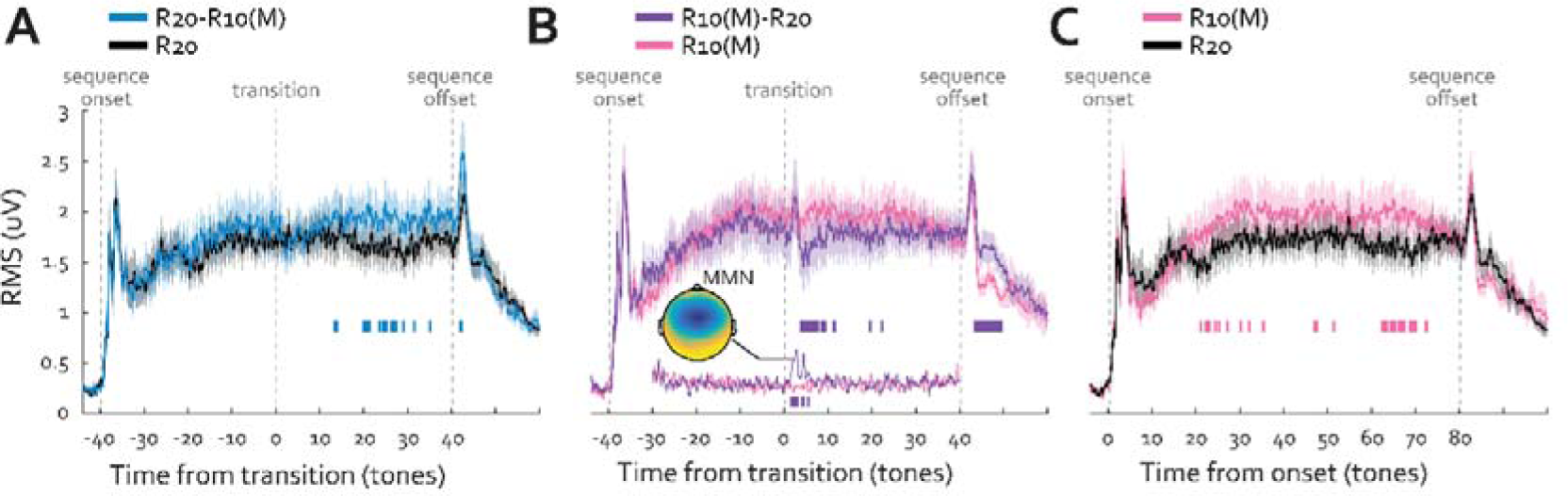
EEG results in Experiment 3 (N=22). EEG responses to R20-R10(M), R10(M) and R10(M)-R20 against their respective controls plotted against time. The shaded area shows 1 SEM (standard error of the mean). Colour-coded horizontal lines at graph bottom indicate time intervals where bootstrap statistics show significant differences between each change condition and its no-change control. (A) The response to R20-R10(M) and its control R20 replicates the pattern observed in Experiment 2A (Figure 4B). (B) The brain response to R10(M)-R20 shows a sharp drop in the sustained response after the transition and then settles at a lower-level post-transition compared with R10(M). The MMN response to the first novel tone in the post-transition sequence can be clearly observed in the 2Hz-high-pass-filtered trace, which is included in the bottom of the plot. The corresponding topography which is consistent with that commonly observed for MMN is also provided. (C) Compared with R20, the brain response to the no-change R10(M) shows a generally higher sustained response, emerging from 21.5 tones after sequence onset.

This suggests that the intrinsic statistics of each stochastic tone distribution, rather than the transition between distributions, may drive the sustained EEG response. Indeed, we observe that the R10(M) no-change condition drives a greater sustained response amplitude than the R20 no-change condition (Figure 5C).

Overall, this pattern is consistent with the hypothesis that the sustained EEG responses reflect a brain state, as opposed to a transition-related response, and moreover that sustained EEG responses track the statistical structure of the sequence. The specific observation of a higher sustained response evoked by R10M than R20 is consistent with the hypothesis from Barascud et al. (27) that the sustained response may track sequence precision: the predictive distribution for R10(M) is expected to be narrower (hence more “precise”) than that for R20.

### Relating D-REX outputs to EEG responses

To determine whether the EEG brain response pattern in naïve distracted listeners can be explained as arising from a process that accumulates distributional statistics, we asked which D-REX model outputs might correspond to the brain responses we observed. As discussed in the introduction, D-REX tracks two key inferred statistics: (1) *Change probability* (the probability that at least one changepoint has occurred); and (2) *Precision* (the reliability of the prediction). In D-REX, precision for multi-gaussian distributions is quantified as the maximum value of the normalised predictive distribution (labelled in Figure 2B).

To simulate the model outputs for the EEG stimuli, we applied the model to the stimuli in Experiment 2. As no behavioural response was acquired in Experiment 2 (passive listening), ideal observer model parameters were used (infinite memory and zero noise). The model was then run on the entire experimental session in the same way it was delivered to each of the 26 participants (i.e., 600 trials, including 100 trials for each transition condition and 300 trials of R20).

Figure 6 shows the model outputs for the “wide-channel” and “multi-channel” priors. Change probability (Figure 6, A & C) grows post-transition, but at rates that differ markedly across conditions. Transition from the full-frequency tone pool (R20) to higher frequency tones only (R10(H)) evoked the steepest increase in change probability, followed by transitions to the middle frequency tone pool (R10(M), and the high and low ‘edge’ tone pool (R10(E)). Precision also grows after the transition, reflecting attunement of the model to the underlying statistics of the sequence (Figure 6, B & D).

**Figure 6.**
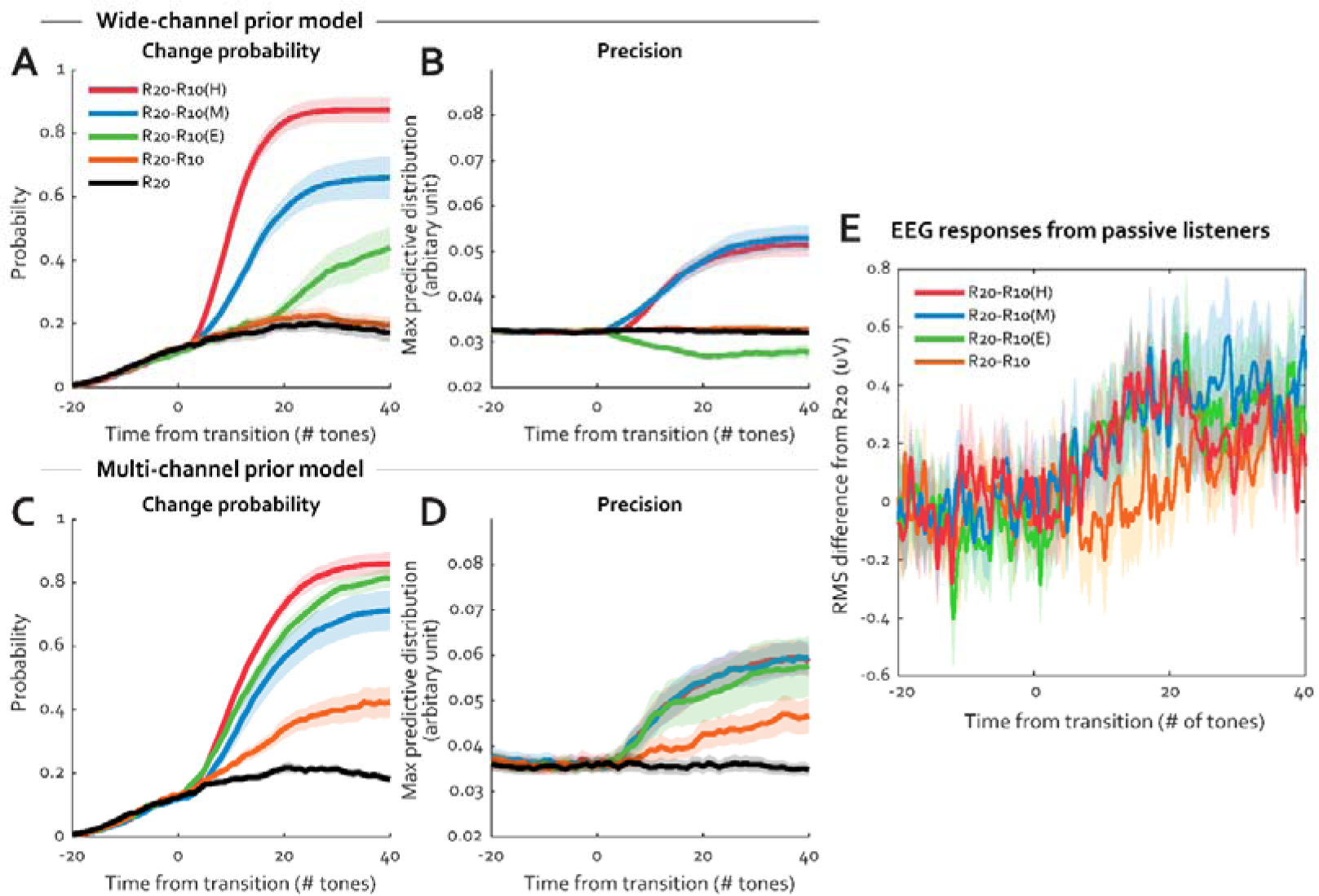
Time-course of modelling outputs and EEG responses from human listeners. Change probability (A) and precision (B) are plotted as a function of time (in number of tones) from the transition for the wide-channel prior model. Outputs of the multi-channel prior model are shown in C and D. For each transition condition, the change probability and precision (see Figure 2B for explanation) were averaged across 100 trials presented to the model, mirroring the evoked response EEG analysis (also based on averaging brain activity across trials). Generally, change probability and precision exhibit a rise, followed by a plateau shortly after the stimulus transition. The specific dynamics reflect the statistics of the stimulus. The unique ‘undershoot’ response observed in the wide-channel prior precision time series for the R20-R10(E) condition (green line in panel B) is a function of both the prior distribution and the underlying distribution of the stimuli. Here, the wide-channel model prediction widens yet further in response to the bimodal stimuli, leading to a reduction in precision from the initial predictive distribution. (E) EEG responses to the four transition conditions (R20-R10(H), R20-R10(M), R20-R10(E) and R20-R10) are overlaid together (previously shown in Figure 4 A-C). Because the responses are taken from two different experiments (Experiment 1A and 1B; see Figure 4) they are normalized by subtracting the relevant R20 baseline activity. The shaded area in A-D is 1 standard deviation of bootstrap of 100 trials presented to the model. The shaded area in E indicates 1 SEM (across participants). All outputs are plotted against time (in #tones) from the transition.

Notably, the wide-channel prior model outputs did not differentiate R20-R10 from R20, either in change probability (Figure 6A) or precision (Figure 6B). The EEG response (Figure 6E) is clearly not consistent with Wide-Channel change probability (Figure 6A) or precision outputs (Figure 6B). By contrast, the “multi-channel” prior model, in particular the “precision” output (Figure 6D) provides a good qualitative match to the EEG responses, with responses to R20-R10(H), R20-R10(M) and R20-R10(E) reaching a similar level after the transition, and with activity in R20-R10 rising at a slower rate.

However, the stimulus set in Experiment 2 is not optimal for adjudicating between multi-channel model predictions because the pattern of results for change probability and precision is quite similar for the three transition stimuli (R20-R10(H), R20-R10(M), R20-R10(E)). In the final experiment we address this issue by employing a different stimulus set for which the model should predict divergent dynamics.

### Automatic brain responses track the precision of unfolding sound sequences

We introduced a new type of transition, going from the full set of 20 to five tones equally distributed across the spectrum (R20-R5), comparing it against our “anchor stimulus” - R20-R10(M) (Figure 7A). R20-R5 was chosen because it has very different underlying stochasticity (see below), which in turn drives different model predictions. In a pilot experiment (see Methods Experiment 4A) we confirmed that R20-5 evokes similar behavioural performance to R20-R10(M).

**Figure 7.**
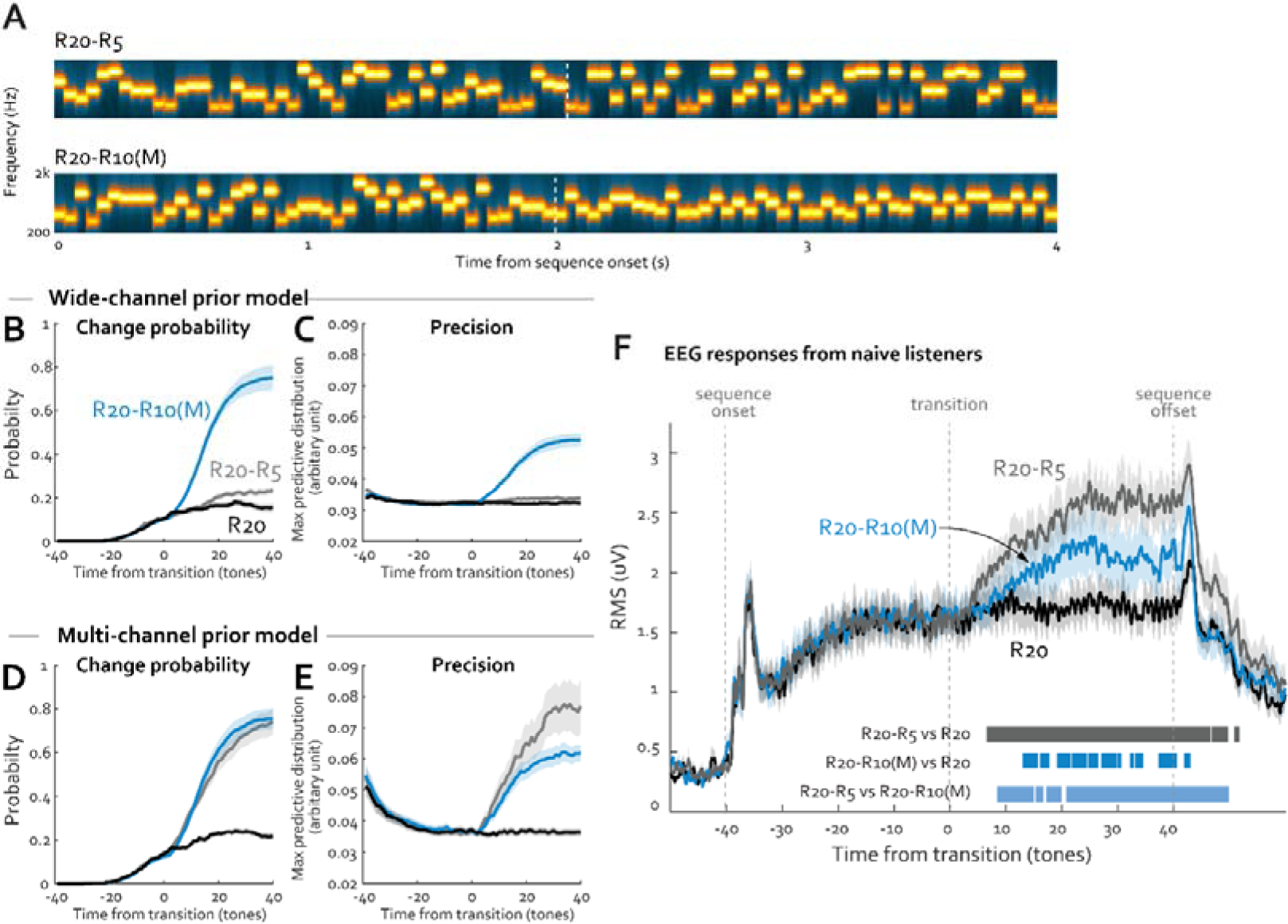
Comparing the brain response and modelling responses to R20-R5 and R20-R10(M) in Experiment 4. (A) Example spectrograms of the two transition conditions used in Experiment. R20-R5 contains a reduction in alphabet size partway through the sequence (from 20 to 5, sampled equally from the full pool). (B) Like Figure 6, A-D, time-courses of model outputs — change probability and precision— are plotted from the transition. The outputs of both wide-channel prior model (B and C) and the multi-channel prior model (D and E) are shown. The outputs were averaged across 100 trials presented to the model. The shaded area in B-E is 1 standard deviation of bootstrap. (F) EEG responses from naïve distracted listeners in Experiment 4B. Plotted are the group mean of RMS over 10 auditory sensitive electrodes. Shaded areas are 1 SEM (standard error of the mean). Colour-coded horizontal lines at graph bottom indicate time intervals where bootstrap statistics show significant differences between each transition condition and R20 (black trace). The bottom horizontal line in light blue indicates intervals where a significant difference between two transition conditions was observed.

Model outputs (change probability and precision against time) for the wide- and multi-channel prior models are plotted in Figure 7, B-E. The wide-channel prior model shows substantially higher change probability (Figure 7B) and precision (Figure 7C) for the R20-R10(M) condition compared to R20-R5. By contrast the multi-channel prior model shows very similar change probability for both transition conditions (Figure 7D), but higher and faster-rising precision *for R20-R5* compared to R20-R10(M) (Figure 7E). This is consistent with the fact that the spectral likelihood distribution for that condition is narrower than for R20-R10(M).

Which model output best predicts passive EEG responses to the same stimuli? In the R20-R10(M) condition, we again find a slow rise in the sustained response (Figure 7F, blue curve), consistent with that seen in Experiments 2A and 3. Critically, the R20-R5 sequence evokes a markedly different response (Figure 7F, grey curve), *one uniquely mirroring the precision pattern predicted by the multi-channel prior model* (Figure 7E). Significantly greater EEG amplitude for R20-R5 versus R20-R10(M) emerged from 422ms after onset and was sustained for the remainder of the trial.

A follow-up, channel-by-channel analysis was conducted to assess the consistency of this effect across scalp loci. We first identified the electrodes (N=40 out of a total of 64) that reliably responded to the transitions (a significant difference between R20-R10(M) and R20 and R20-R5 and R20 in the interval 1-2 seconds post-transition; see *Methods - EEG channel-by-channel analysis*). For each of the transition-sensitive electrodes, we then computed the amplitude difference between R20-R5 and R20-R10(M) at 1-2 seconds post transition. These values are illustrated in Figure 8. This analysis confirmed that no electrode exhibited R10M>R5, providing further evidence against the “wide-channel” model. The vast majority of the transition-sensitive electrodes exhibited a larger response to R5 relative to R10(M), indicative of precision tracking, and in-line with the RMS analysis above.

**Figure 8.**
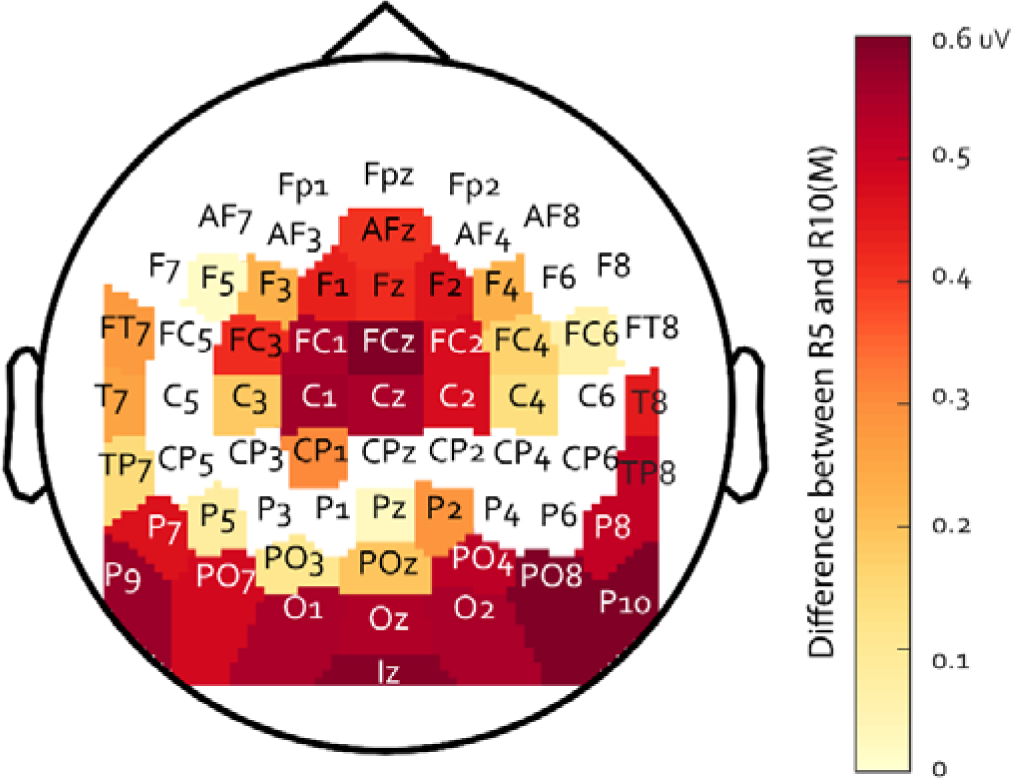
Channel-by-channel analysis of responses to R20-R5 and R20-R10(M) in Experiment 4. Plotted is the mean difference during the 1-2 second post-transition period between these conditions in each electrode. A higher positive value signifies a higher amplitude response to R5 compared to R10(M).

The close similarity between the D-REX Multi-Channel precision output and these EEG data suggests that the passively listening brain utilises a within-channel listening strategy, and that the brain responses driving the EEG sustained response reflect the precision of the internal model of the sound sequence constructed during listening.

## Discussion

Listeners automatically track the statistics of unfolding sounds even when these are not relevant to behaviour. Transitions from R20 (uniform sampling from the full frequency pool) to various constrained sampling regimes drove an increase of the sustained EEG response. This was observed to a range of distributional statistics, and even in situations (R20-R10) where behavioural detection of these transitions in Experiment 1 was at floor. Similar EEG responses were previously reported for deterministic patterns (17,27–31). That they are also seen for stochastic transitions supports the hypothesis that EEG sustained response modulation reflects a general process of belief updating.

The EEG data were compared to a Bayesian predictive inference model fit to the acoustic scenes experienced by listeners. This comparison showed that EEG responses were most closely associated with coding *precision*, a representation of the uncertainty associated with the current predictive distribution.

### Listeners are sensitive to stochastic regularities

Accumulating work demonstrates that the human brain is tuned to the statistical properties of the stochastic ongoing acoustic environment (3,7,41–45). By modelling brain responses to two-tone sequences of different base probabilities, Rubin et al. (45) showed that trial-wise neural responses in auditory cortex are well explained by the probability of occurrence of each tone frequency, calculated from the recent history of the sequence. The models that best fit neural responses tracked a relatively long stimulus history (∼10 tones) but encoded a coarse representation of stimulus properties, based on a small set of 0-th order statistics. In line with this conclusion, Garrido et al. (41) (also see 45,46) demonstrated that MEG responses to probe (outlier) tones are sensitive to the statistical context (mean and variance of frequency) of randomly generated tone-pip sequences. Larger responses occurred to the same probe tone when presented in a context with low, as opposed to high, frequency variance. However, through its emphasis on the effect of various statistical contexts on the processing of *deviant* tones, this work has provided largely indirect insight into the brain and perceptual mechanisms through which listeners track the statistics of stochastic auditory environments.

More recently, several investigations have focused on the factors that affect change detection in stochastic sound sequences. Boubenec et al. (6) used a broadband ‘tone cloud’ stimulus composed of tone-pips randomly drawn from a spectral probability distribution. They demonstrated that listeners’ change detection accuracy depended on the spectral properties of the changed distribution. Change- and decision-evoked EEG responses were identified in centro-parietal electrodes, consistent with other evidence integration tasks (48–50). Using fractal sequences, Skerritt-Davis & Elhilali (3,7) found that listeners’ change detection and surprisal-associated brain responses were consistent with an observer model that tracks the higher-order statistics of temporal covariance structures in the sequences.

Whereas these previous studies used active monitoring of changes-points in auditory sequences, the present study demonstrates automatic statistical tracking during passive listening. The brain responses implicated in belief updating originated from central scalp locations, likely reflecting auditory cortical activity. The electrodes exhibiting the sustained response modulation were also those that responded most strongly to auditory onsets, suggesting that auditory cortex is involved in tracking these statistics. This contrasts with Boubenec et al. (6) where evidence accumulation-activity was concentrated in centro-parietal electrodes. The difference between our findings and those in (6) could potentially be attributed to task-related factors. In our study, the observed responses reflect automatic tracking, in contrast to Boubenec et al. (6) where participants actively monitored for sound changes, engaging explicit “evidence accumulation” mechanisms.

### Bayesian tracking of rapidly unfolding sounds

We used a Bayesian model of statistical tracking (“dynamic regularity extraction”, D-REX (37)) to constrain our interpretation of EEG dynamics. A detailed account of the model is presented elsewhere (37), see also (3,7). D-REX is similar to frameworks widely used to model various aspects of human behaviour under uncertainty (1–4,7,16,38,51,52). Specifically, the model is based on key assumptions in predictive coding: the brain (a) builds beliefs by collecting statistical estimates over time; and (b) simultaneously maintains multiple hypotheses regarding the stimulation context it has just experienced. The latter assumption incorporates a focus on detecting changes in the underlying generative distribution, often referred to as “change point detection.” Each sequentially encountered stimulus is used to update these hypotheses in an “*observe-predict-update”* loop (53–55).

Prior work in decision-making used a modelling approach similar to D-REX to show that listeners track change point probability, and use it to optimize their prediction of future variance (1,2,16). In the context of auditory processing, D-REX has been used to model a variety of perceptual phenomena related to understanding the extent to which auditory information is represented in memory (3,7,37). In tasks that required listeners to monitor for changes in rapid sound sequences, the model successfully fit performance. The model results suggested that listeners track marginal mean variance, and covariance (temporal dependency) between successive elements. The free model parameters: *memory* (maximum context window) and *observation noise* accounted for individual variability in reaction time and sensitivity. Similar effects were seen in our behavioural experiment (Experiment 1). We observed high correlations between model performance and individual subject behaviour, in particular for the memory parameter which suggested that listeners use a context of 20-80 tones to inform their decision about change occurrence.

### EEG responses in naïve listeners reflect within-channel precision tracking

To explain the dynamics of EEG responses, we focused on two key instantaneous outputs of the model: 1), its beliefs about *the probability that change point has occurred*, i.e., that the generative distribution underlying the stimulus sequence has changed; and 2), *the precision of its predictive representation*, reflecting the accuracy (or conversely ‘expected uncertainty’; O’Reilly, 2013) with which future inputs can be predicted. We also evaluated two potential listening strategies by initiating the model with two types of priors: either a single Gaussian (*Wide Single-channel prior*), reflecting the hypothesis that listeners initially track a single wide-band spectral distribution; or a multimodal Gaussian (*Multi-channel prior*), where listeners employ a within-frequency-channel listening strategy, where several concurrent spectral distributions are initially tracked.

Relating the EEG responses to outputs of the D-REX model demonstrated that the sustained response pattern was consistent with the “multi-channel prior” model. This is in line with many existing demonstrations (56–58) that the auditory system’s “default mode” is to process sound via multiple separate spectral channels. Interestingly, it appears that behavioural performance (Experiment 1, Figure 3) was better modelled with the “wide-channel prior” model, hinting at a potential distinction between active statistical tracking and passive listening, an initial finding which should be followed up in future work.

Earlier observations in the context of deterministic sequences showed that the sustained response is modulated by sequence precision: Regular patterns consistently evoke larger sustained EEG responses than do random patterns (27–31). In addition, more predictable random patterns (i.e., smaller alphabet size) evoke higher sustained EEG responses than less predictable random patterns (27). Another study (17) also reported similar effects in the context of artificial “scenes” that were populated by temporally regular vs. random streams. Extending these observations to stochastic patterns, we show that the sustained response recorded in naïve distracted listeners reflects a brain state that varies systematically with sequence precision. This is consistent with precision weighting being an optimal evidence integration strategy (8), and establishes the sustained M/EEG response as a potential neural signature for precision tracking.

The ability to accurately estimate precision allows an organism to operate optimally under different levels of environmental volatility by reweighing signals according to their inferred reliability (8,59). This increases the salience of deviants in a high precision context, indicative of a genuine change in the environment. Generally, high precision signifies a stable and predictable environment, which organisms tend to favour (8). From a survival standpoint, it likely triggers a sense of relative safety, activating the parasympathetic nervous system and facilitating a “rest-and-digest” mode (60). This state may be reflected by the large shift in sustained response we observe.

That EEG responses do *not* reflect change probability tracking might be somewhat surprising given the key importance of sensitivity to environmental change to perception and survival. However, this finding is consistent with observations from Boubenec et al. (6) who also did not find any evidence for change-point detection-related activity in auditory cortex. One possibility is that change probability *is* being tracked in auditory cortex but in a way that is not reflected in the EEG activity we analysed.

### The neural correlates of the EEG sustained response

The neural-circuit underpinnings of the sustained-response modulation effects remain elusive. According to one proposal, precision is encoded by gain in superficial pyramidal cells in sensory cortex (12,34), potentially via neuromodulation. In particular, tonic Acetylcholine (ACh) has been shown to be modulated by environmental uncertainty (61–63).

An alternative, though not mutually exclusive, account links precision tracking to inhibitory mechanisms. Growing evidence from animal electrophysiology and modelling (albeit using very simple sounds) demonstrates that inhibitory plasticity plays a critical role in shaping brain responses to repeating stimuli. Specifically, increased inhibition onto excitatory neurons tuned to familiar (repeating/predictable) stimuli might constitute a cortical mechanism for precision tracking (64–68). Under this hypothesis the modulations in the sustained response may be reflecting increased inhibitory activity. Indirect evidence from recent dynamic causal modelling (DCM) work has also suggested that encoding of precision is attributable to inhibitory mechanisms (26).

A specific role for inhibition in controlling responses to predictable sensory stimuli is also consistent with behavioural results showing that rather than attracting attention (which would be expected from an excitatory effect), high-precision patterns are easier to ignore (28), and are associated with reduced arousal (60). M/EEG (or BOLD) cannot straightforwardly resolve inhibitory from excitatory activity. Progress in testing this account therefore depends on targeted cellular-level or pharmacological investigations.

## Materials and Methods

### Ethics Statement

Experimental procedures were approved by the research ethics committee of University College London, and written informed consent was obtained from each participant. Participants were paid for their participation.

### Stimuli

The stimuli were constructed by selecting tone-pips (50ms long, gated with 5ms raised-cosine ramps) from a pool of 20 frequencies (‘alphabet’; 222–2000Hz, ∼12.2% steps or ∼2 semitones, as in Barascud et al. (27)). The frequency steps provide sufficient spacing to allow differentiation, while the short unit ensures that listeners cannot explicitly track the sequence (69–71).

The basic stimuli were random sequences constructed by sampling frequencies uniformly at random from the full pool (“alphabet” =20). This condition is referred to as “R20” (Figure 1, top). Transition stimuli were created by altering the sampling regime partway through the sequence.

The full pool of 20 frequencies was divided into four logarithmically equally sized frequency bands, each containing five frequency values. Different conditions were created by reducing the *alphabet* to 10 or 5 frequencies, and manipulating how these are selected. Spectrograms of example stimuli can be found in Figure 1. Sound examples can be found online [https://github.com/sijiazhao/share_ranran].

In **R20-R10**, the fully random (R20) sequence transitioned into a sequence that contained 10 frequencies (R10) uniformly sampled from across the full pool (2 from each band).

a. **R20-R10(H)** contained a transition into a sequence comprising 10 frequencies sampled from the two highest bands (707Hz to 2000Hz).
b. **R20-R10(L)** contained a transition into a sequence comprising 10 frequencies sampled from the two lowest bands (222Hz to 630Hz).
c. **R20-R10(M)** contained a transition into a sequence comprising 10 frequencies sampled from the two middle bands (397Hz to 1122Hz).
d. **R20-R10(E)** contained a transition into a sequence comprising 10 frequencies sampled from the highest (1259Hz to 2000Hz) and lowest (222Hz to 354Hz) bands. In Experiment 4 we also used
e. **R20-R5** which contained a transition into a sequence comprising 5 frequencies uniformly sampled from across the full pool (1 from each band; 1 selected at random from the remaining values). (Figure 7A)

All stimuli were generated anew for each trial in each subject. In the behavioural experiments (Experiment 1 and Experiment 4A), the transition time was jittered between 40-50 tones after sequence onset. In all EEG experiments (Experiments 2A, 2B, 3 and 4B), the sequence length for each trial was fixed to 80 tones and the change position was fixed to the 41st tone.

### Experiment 1

#### Participants

Ten participants (10 females; aged 19–24, average 21.6) took part in the experiment.

#### Stimuli

The stimulus set included R20 (control) and five transition conditions: R20-R10(H), R20-R10(L), R20-R10(M), R20-R10(E), and R20-R10. Each condition was presented in a separate block (20 transition conditions plus 20 R20 controls; block order was randomized across participants). Transition times were jittered by assigning a random length to the pre-transition segment; the transition time was therefore unpredictable. Trials were 80–90 tone-pips long, lasting 4–4.5 seconds.

To make this obvious change unpredictable, the transition time and sequence length were also jittered in exactly the same way as for the experimental transition conditions. The inter-trial interval (ITI) was randomised between 1 and 1.5 seconds.

The stimuli were delivered to the participants’ ears by Sennheiser HD558 headphones (Sennheiser, Germany) via a UA-33 sound card (Roland Corporation) at a comfortable listening level, self-adjusted by each participant. Stimulus presentation and response recording were controlled with the Psychtoolbox package (Psychophysics Toolbox Version 3)(72) in MATLAB (The MathWorks, Inc.).

Participants were tested in a darkened, acoustically shielded room (IAC triple-walled sound-attenuating booth). Each experimental session lasted about 1 hour and began with a short practice session, followed by the main experiment. Participants were instructed to fixate their gaze on a white cross at the centre of the computer screen while listening to the stimuli and to respond by pressing a keyboard button as soon as they detected the transitions. Feedback (in the form of a tick or a cross) was provided after each trial. Each block lasted about 6 minutes and participants were allowed short breaks between blocks.

### Experiment 2 - EEG

#### Participants

**Experiment 2A:** Data from 26 naïve participants (18 females; aged 21–38; average 24.1; 2 left-handed) are reported. No participant was excluded in Experiment 2A.

**Experiment 2B:** Data from 26 naive participants (17 females; aged 20–32, average 23.8, all right-handed) are presented. One additional participant was rejected due to excessive missing data (one block of data failed to record).

All participants reported normal hearing with no history of neurological disorders. None of the participants was a professional musician or known to possess absolute pitch.

#### Stimuli

**Experiment 2A:** The stimulus set included R20-R10(H), R20-R10(M), R20-R10(E) and their

non-change control R20. Overall, 600 stimuli (100 each of R20-R10(H), R20-R10(M), R20-R10(E), and 300 of R20) were presented to the listeners in a random order.

**Experiment 2B:** The stimulus set consisted of two stimulus conditions: R20-R10 and its no-change control R20. In total 200 stimuli (100 each of R20-R10 and R20) were presented in random order.

All stimuli were 4 seconds long (80 tones) with the transition occurring at 2 seconds post onset (at the 41st tone-pip). The inter-trial interval (ITI) varied between 1.5 and 2 seconds. Stimuli were generated offline in MATLAB and presented binaurally with the Psychophysics Toolbox (72,73) using EarTone in-ear earphones at a comfortable listening level of ∼60 dB SPL self-adjusted by each participant.

#### Procedure

After the placement of the electrodes, participants were seated in front of a computer monitor and instructed to watch a silent subtitled documentary throughout data collection. The monitor was adjusted to present the subtitles at eye level. The experiment was organized into 10-minute blocks (6 blocks in Experiment 2A; 2 blocks in Experiment 2B). Short breaks were provided between blocks, but participants were asked to remain still in the breaks.

EEG was recorded through a 64 channel Biosemi system (Biosemi Active Two AD-box ADC-17, Biosemi, Netherlands) with a 64-electrode cap, referenced to the CMS-DRL ground, which functions as a feedback loop driving the average potential across the montage as close as possible to amplifier zero. Acquisition was continuous with a sampling rate of 2048 Hz.

### Experiment 3 - EEG

#### Participants

All participants from Experiment 2B also took part in this experiment; however, data from four participants were excluded due to excessive missing data (one block of data failed to record). Data from 22 participants are presented (13 females; aged 21–32, average 24, all right-handed). All participants, first completed Experiment 3, followed by Experiment 2B.

#### Stimuli

This experiment contained four stimulus conditions: R20-R10(M) and its control R20, and R10(M)-R20 and its control R10(M). Each were presented 100 times in random order. The R10(M)-R20 stimuli were constrained such that the first tone after the transition was novel (i.e., not contained in the preceding R10M sequence). This ensured that statistically, the transition occurred at the same time in all stimuli. The experiment was organized into four 10-minute blocks. Short breaks were provided between blocks, but participants were asked to remain still in the breaks.

### Experiment 4A – Behavioural Pilot

#### Participants

Five new participants (5 females; aged 21–23, average 22) participated in the pilot.

#### Stimuli and Procedure

The experiment contained 80 trials: 20 trials of R20-R5, 20 trials of R20-R10(M) and 40 trials of R20. All were randomly mixed. Otherwise, the procedure was identical to Experiment 1.

### Experiment 4B – EEG

#### Participants

28 new participants (18 females, aged between 22 and 34, average 24.1, 2 left-handed) participated this experiment. Four participants were excluded, three due to excessive missing data (one block of data failed to record) and one due to faulty earphones. Data from 24 participants (15 females, aged 22-34 average 24.2; 1 left-handed) are presented.

#### Stimuli

This experiment contained three conditions: R20-R5, R20-R10(M) and their common control R20. Transition times were as in Experiment 2, above. R5 sequences were created by randomly retaining 5 frequencies from the full pool – 1 from each band and an additional value randomly selected from across all bands. Overall, 600 stimuli (150 R20-R5, 150 R20-R10(M) and 300 of R20) were presented to the listeners in a random order. The experiment was organized into six 10-minute blocks. Short breaks were provided between blocks, but participants were asked to remain still in the breaks.

### Data Analysis

#### EEG Data Processing

The EEG analysis was performed using the Fieldtrip toolbox (74) in MATLAB (2015b, The MathWorks, Inc., Natick, Massachusetts, United States).

Data were first smoothed by convolution with a square window of size 1/50 Hz to suppress 50 Hz and harmonics, and then low-passed at 30 Hz (two-pass, fifth-order Butterworth filter). Next, the data were downsampled to 256 Hz, and 5500 ms epochs (from 500 ms pre-onset to 1000 ms post-offset) were extracted for each trial. In Experiments 2A, 3, and 4B, the number of R20 trials was larger than that of the other conditions. To equate SNR between conditions, the number of trials contributing to the estimation of R20 activity was reduced to match the trial number of the other transition conditions by random selection. Data were then detrended using the NoiseTools toolbox (75) and average-referenced across the full set of channels.

De-noising source separation (DSS) analysis was applied to maximize reproducibility across trials (76,77). In Experiments 2 and 4B DSS was applied on collapsed data (across conditions). In Experiment 3 DSS was run separately for each condition. The rationale was that the transition condition R10(M)-R20 might evoke a brain response (e.g., MMN) which might not share the components with other conditions.

For each participant, the first five DSS components (i.e., the five most reproducible components) were selected and projected back into sensor space. Trials which deviated from the mean by more than twice the standard deviation were automatically flagged as outliers (normally less than 10%) and discarded from further analyses. Subsequently, the data were averaged across trials.

To identify additional (fast) activity, which may potentially be masked by the slow changes, the same analysis as described above was also performed on 2 Hz high-pass filtered data (two-pass, sixth-order Butterworth filter).

The root-mean-square (RMS) of the measured current at each time point was calculated over a subset of 10 channels (5 central, 5 occipital: FCz, Fz, Cz, FC1, FC2, P9, P10, Iz, O1 and O2) for each participant and condition and used as a measure of brain activation over time. These channels were selected because they were the most activated during the N1 onset response (around 100 ms after stimulus onset; see Figure 4A) in the grand average data (computed by collapsing across conditions) and thus considered to best capture auditory activity.

#### EEG Timeseries Statistical Analysis

For illustration purposes, group-RMS (RMS of individual RMSs) is shown, but statistical analysis is always conducted across participants. The difference between the (squared) RMS waveforms of each transition condition and R20 was calculated for each participant, and subjected to bootstrap re-sampling (5000 interactions; balanced) (78). The difference was deemed significant if the proportion of bootstrap iterations that fell above or below zero was more than 95% (i.e., p < 0.05) for 10 or more consecutive samples (i.e., > 39 ms). The significant intervals determined in this way are indicated as horizontal lines, below the EEG trace. The time at which the first significant sample is observed is considered the earliest time point at which the brain has detected that the statistics of the stimulus have changed.

The threshold for the number of consecutive samples (10) was determined by comparing activity within the initial 2000 ms across conditions. Stimuli are identical during this period (R20); therefore, any significant differences are fully attributable to noise. We estimated the largest number of consecutive samples to show a significant effect in that interval by rerunning the analysis on the −500 to 2000ms interval 1000 times and taking the longest obtained “spurious” significant interval as the threshold for the statistical analysis.

#### EEG channel-by-channel analysis (Experiment 4)

This analysis sought to assess the consistency of the EEG effect R5 > R10(M) across various EEG electrodes in Experiment 4B. Initially, we identified the electrodes sensitive to transitions, where both R20-R5 and R20-R10(M) exhibited significant deviations from R20 within the 1-2 second post-transition timeframe. This selection process yielded 40 out of the 64 available electrodes.

For each of the ‘transition-sensitive’ electrodes, we conducted baseline correction on the absolute time series of R20-R5 and R20-R10(M) using a 0.5-second pre-transition interval, followed by subtraction. The mean difference during the 1-2 second post-transition period was computed. A higher positive value signifies a more pronounced response to R5 compared to R10(M). These values are depicted in Figure 8 with colour coding.

In Figure 8, the majority of the transition-sensitive electrodes display large positive values. Notably, none of the electrodes exhibited negative values, indicating that none of the channels demonstrated a response pattern where R5 < R10(M).

#### Behavioural Analysis

The dependent measure was *d’* (79). For the purposes of computing the *d’* score, transition trials in which the participant responded after the nominal transition time were considered as hits. Responses to no change trials, or before the onset of the transition were labelled as false positives.

Statistical analysis was performed using SPSS. In all analyses of variance (ANOVA), the Greenhouse-Geisser correction was applied when the assumption of sphericity was violated (80). The _α_ level was set a priori to 0.05.

### Computational Modelling: D-REX model

An auditory perceptual model - Dynamic Regularity Extraction (D-REX) - was used to capture and predict the unfolding statistics within sequential experimental stimuli (7). The model is an extension of a general Bayesian Online Changepoint Detection model (81) which uses Bayesian inference to perform sequential predictions given previous observations in the presence of unknown changepoints.

The model operates on a basic assumption that all observations are from a D-dimensional Gaussian distribution θ. At time t, this distribution can be estimated by sufficient statistics 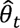 using the *r_t_*, previous run of observations : 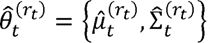, where 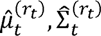are the sample mean and sample covariance, respectively, estimated from *r_t_* (82).

Two perceptual parameters are also used in this model: 1) memory (m), which limits the extent of previous observations *r_t_* used to estimate sufficient statistics for generating predictions; and 2) observation noise (n), which represents limitations on perceptual fidelity by adding independent Gaussian noise to the estimation of the predictive distribution. Hence, the predictive distribution at time t with the previous observations *r_t_* is:

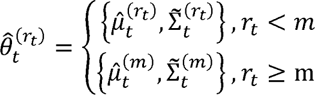

where 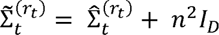 is the sample covariance with added observation noise n, and I is the D-dimensional identity matrix. The sufficient statistics underlying the predictive probability track changes in a multivariate Gaussian distribution in line with previous work (37).Two versions of the model were compared: 1) A wide-channel model which uses a single Gaussian distribution as initial prior; and 2) a multi-channel model which uses a multimodal Gaussian prior. In the latter case, the prior distribution uses 5 channels spread evenly in log-frequency (between −16 to 16 semitones relative to the centre frequency of the stimuli). The choice of five channels was motivated by the intention to mitigate any potential bias in the results towards the frequency bands employed in stimulus generation.

The model is implemented in MATLAB and its code is available online https://github.com/JHU-LCAP/DREX-model. The model implementation is overall similar to that described in detail in a previous study (37) with the following modifications: First, all inputs are assumed to be independent (no temporal covariance). Second, the hazard function, which sets the prior probability of change occurrence is set to 0.01. This means that the model is conservative in its expectation of change. For behavioural experiments, an individual D-REX model was matched to each subject by performing a grid search on the best model parameters (memory m and noise n) that replicate the subject’s responses. For EEG experiments, a single ideal observer model was used with infinite memory and zero noise.

## Acknowledgements

This work was supported by a BBSRC project grant to MC and ONR grant (N00014-23-1-2050 to ME. We are grateful to Daniel Yon and Clare Press for comments and discussion.

## References

1. Nassar MR, Wilson RC, Heasly B, Gold JI. An Approximately Bayesian Delta-Rule Model Explains the Dynamics of Belief Updating in a Changing Environment. J Neurosci. 2010 Sep 15;30(37):12366–78.

2. Nassar MR, Rumsey KM, Wilson RC, Parikh K, Heasly B, Gold JI. Rational regulation of learning dynamics by pupil-linked arousal systems. Nat Neurosci. 2012 Jul;15(7):1040–6.

3. Skerritt-Davis B, Elhilali M. Neural Encoding of Auditory Statistics. J Neurosci. 2021 Aug 4;41(31):6726–39.

4. Wilson RC, Nassar MR, Gold JI. A Mixture of Delta-Rules Approximation to Bayesian Inference in Change-Point Problems. PLOS Computational Biology. 2013 Jul 25;9(7):e1003150.

5. Glaze CM, Kable JW, Gold JI. Normative evidence accumulation in unpredictable environments. Behrens T, editor. eLife. 2015 Aug 31;4:e08825.

6. Boubenec Y, Lawlor J, Górska U, Shamma S, Englitz B. Detecting changes in dynamic and complex acoustic environments. Behrens TE, editor. eLife. 2017 Mar 6;6:e24910.

7. Skerritt-Davis B, Elhilali M. Detecting change in stochastic sound sequences. PLOS Computational Biology. 2018 May 29;14(5):e1006162.

8. Yon D, Frith CD. Precision and the Bayesian brain. Current Biology. 2021 Sep 13;31(17):R1026–32.

9. Lawson RP, Bisby J, Nord CL, Burgess N, Rees G. The Computational, Pharmacological, and Physiological Determinants of Sensory Learning under Uncertainty. Curr Biol. 2021 Jan 11;31(1):163–172.e4.

10. Heilbron M, Chait M. Great Expectations: Is there Evidence for Predictive Coding in Auditory Cortex? Neuroscience. 2018 Oct 1;389:54–73.

11. Kanai R, Komura Y, Shipp S, Friston K. Cerebral hierarchies: predictive processing, precision and the pulvinar. Phil Trans R Soc B. 2015 May 19;370(1668):20140169.

12. Feldman H, Friston KJ. Attention, Uncertainty, and Free-Energy. Frontiers in Human Neuroscience [Internet]. 2010 Dec 2 [cited 2017 Jan 25];4. Available from: http://www.ncbi.nlm.nih.gov/pmc/articles/PMC3001758/

13. Weissbart H, Kandylaki KD, Reichenbach T. Cortical Tracking of Surprisal during Continuous Speech Comprehension. Journal of Cognitive Neuroscience. 2020 Jan 1;32(1):155–66.

14. Friston KJ. Precision Psychiatry. Biological Psychiatry: Cognitive Neuroscience and Neuroimaging. 2017 Nov 1;2(8):640–3.

15. Fitzgerald J, Johnson K, Kehoe E, Bokde ALW, Garavan H, Gallagher L, et al. Disrupted Functional Connectivity in Dorsal and Ventral Attention Networks During Attention Orienting in Autism Spectrum Disorders. Autism Research. 2015;8(2):136–52.

16. Krishnamurthy K, Nassar MR, Sarode S, Gold JI. Arousal-related adjustments of perceptual biases optimize perception in dynamic environments. Nat Hum Behav. 2017;1:0107.

17. Sohoglu E, Chait M. Detecting and representing predictable structure during auditory scene analysis. eLife. 2016 Sep 7;5:e19113.

18. Barczak A, O’Connell MN, McGinnis T, Ross D, Mowery T, Falchier A, et al. Top-down, contextual entrainment of neuronal oscillations in the auditory thalamocortical circuit. PNAS. 2018 Aug 7;115(32):E7605–14.

19. Asokan MM, Williamson RS, Hancock KE, Polley DB. Inverted central auditory hierarchies for encoding local intervals and global temporal patterns. Current Biology. 2021 Apr;31(8):1762–1770.e4.

20. Demarchi G, Sanchez G, Weisz N. Automatic and feature-specific prediction-related neural activity in the human auditory system. Nature Communications [Internet]. 2019 Dec [cited 2019 Aug 6];10(1). Available from: http://www.nature.com/articles/s41467-019-11440-1

21. Garrido MI, Dolan RJ, Sahani M. Surprise Leads to Noisier Perceptual Decisions. i-Perception. 2011 Feb 1;2(2):112–20.

22. Hsu YF, Hämäläinen JA. Both contextual regularity and selective attention affect the reduction of precision-weighted prediction errors but in distinct manners. Psychophysiology. 2021 Mar;58(3):e13753.

23. Sedley W, Gander PE, Kumar S, Kovach CK, Oya H, Kawasaki H, et al. Neural signatures of perceptual inference. eLife. 5:e11476.

24. Hsu YF, Waszak F, Hämäläinen JA. Prior Precision Modulates the Minimization of Auditory Prediction Error. Frontiers in Human Neuroscience [Internet]. 2019 [cited 2023 Oct 17];13. Available from: https://www.frontiersin.org/articles/10.3389/fnhum.2019.00030

25. SanMiguel I, Costa-Faidella J, Lugo ZR, Vilella E, Escera C. Standard Tone Stability as a Manipulation of Precision in the Oddball Paradigm: Modulation of Prediction Error Responses to Fixed-Probability Deviants. Frontiers in Human Neuroscience [Internet]. 2021 [cited 2023 Oct 17];15. Available from: https://www.frontiersin.org/articles/10.3389/fnhum.2021.734200

26. Lecaignard F, Bertrand O, Caclin A, Mattout J. Neurocomputational Underpinnings of Expected Surprise. J Neurosci. 2022 Jan 19;42(3):474–86.

27. Barascud N, Pearce MT, Griffiths TD, Friston KJ, Chait M. Brain responses in humans reveal ideal observer-like sensitivity to complex acoustic patterns. Proceedings of the National Academy of Sciences. 2016 Feb 2;113(5):E616–25.

28. Southwell R, Baumann A, Gal C, Barascud N, Friston K, Chait M. Is predictability salient? A study of attentional capture by auditory patterns. Phil Trans R Soc B. 2017;372(1714):20160105.

29. Southwell R, Chait M. Enhanced deviant responses in patterned relative to random sound sequences. Cortex. 2018 Dec 1;109:92–103.

30. Herrmann B, Araz K, Johnsrude IS. Sustained neural activity correlates with rapid perceptual learning of auditory patterns. NeuroImage. 2021 Sep 1;238:118238.

31. Herrmann B, Johnsrude IS. Neural Signatures of the Processing of Temporal Patterns in Sound. J Neurosci. 2018 Jun 13;38(24):5466–77.

32. Megela AL, Teyler TJ. Habituation and the human evoked potential. J Comp Physiol Psychol. 1979 Dec;93(6):1154–70.

33. Pérez-González D, Malmierca MS. Adaptation in the auditory system: an overview. Front Integr Neurosci [Internet]. 2014 Feb 21 [cited 2016 Mar 9];8. Available from: http://www.ncbi.nlm.nih.gov/pmc/articles/PMC3931124/

34. Auksztulewicz R, Barascud N, Cooray G, Nobre AC, Chait M, Friston K. The cumulative effects of predictability on synaptic gain in the auditory processing stream. J Neurosci. 2017 Jun 12;0291–17.

35. Harrison AW, Mannion DJ, Jack BN, Griffiths O, Hughes G, Whitford TJ. Sensory attenuation is modulated by the contrasting effects of predictability and control. NeuroImage. 2021 Aug 15;237:118103.

36. Andrillon T, Kouider S, Agus T, Pressnitzer D. Perceptual learning of acoustic noise generates memory-evoked potentials. Curr Biol. 2015 Nov 2;25(21):2823–9.

37. Skerritt-Davis B, Elhilali M. Computational framework for investigating predictive processing in auditory perception. Journal of Neuroscience Methods. 2021 Aug 1;360:109177.

38. Behrens TEJ, Woolrich MW, Walton ME, Rushworth MFS. Learning the value of information in an uncertain world. Nature Neuroscience. 2007 Sep;10(9):1214–21.

39. Payzan-LeNestour E, Bossaerts P. Risk, Unexpected Uncertainty, and Estimation Uncertainty: Bayesian Learning in Unstable Settings. PLOS Computational Biology. 2011 Jan 20;7(1):e1001048.

40. Gold JI, Stocker AA. Visual Decision-Making in an Uncertain and Dynamic World. Annual Review of Vision Science. 2017 Sep 15;3(1):227–50.

41. Garrido MI, Sahani M, Dolan RJ. Outlier Responses Reflect Sensitivity to Statistical Structure in the Human Brain. PLoS Comput Biol. 2013 Mar 28;9(3):e1002999.

42. Winkler I, Paavilainen P, Alho K, Reinikainen K, Sams M, Naatanen R. The Effect of Small Variation of the Frequent Auditory Stimulus on the Event-Related Brain Potential to the Infrequent Stimulus. Psychophysiology. 1990 Mar 1;27(2):228–35.

43. Overath T, Cusack R, Kumar S, von Kriegstein K, Warren JD, Grube M, et al. An Information Theoretic Characterisation of Auditory Encoding. PLoS Biol. 2007 Oct 23;5(11):e288.

44. Furl N, Kumar S, Alter K, Durrant S, Shawe-Taylor J, Griffiths TD. Neural prediction of higher-order auditory sequence statistics. NeuroImage. 2011 Feb 1;54(3):2267–77.

45. Rubin J, Ulanovsky N, Nelken I, Tishby N. The Representation of Prediction Error in Auditory Cortex. PLOS Computational Biology. 2016 Aug 4;12(8):e1005058.

46. Daikhin L, Ahissar M. Responses to deviants are modulated by subthreshold variability of the standard. Psychophysiology. 2012 Jan 1;49(1):31–42.

47. Herrmann B, Parthasarathy A, Han EX, Obleser J, Bartlett EL. Sensitivity of rat inferior colliculus neurons to frequency distributions. Journal of Neurophysiology. 2015 Nov;114(5):2941–54.

48. O’Connell RG, Dockree PM, Kelly SP. A supramodal accumulation-to-bound signal that determines perceptual decisions in humans. Nat Neurosci. 2012 Dec;15(12):1729–35.

49. Kelly SP, O’Connell RG. Internal and External Influences on the Rate of Sensory Evidence Accumulation in the Human Brain. J Neurosci. 2013 Dec 11;33(50):19434–41.

50. Yao JD, Gimoto J, Constantinople CM, Sanes DH. Parietal Cortex Is Required for the Integration of Acoustic Evidence. Current Biology. 2020 Sep 7;30(17):3293–3303.e4.

51. Daw ND, O’Doherty JP, Dayan P, Seymour B, Dolan RJ. Cortical substrates for exploratory decisions in humans. Nature. 2006 Jun 15;441(7095):876–9.

52. Wilson R, Niv Y. Inferring Relevance in a Changing World. Frontiers in Human Neuroscience [Internet]. 2012 [cited 2023 Oct 17];5. Available from: https://www.frontiersin.org/articles/10.3389/fnhum.2011.00189

53. Knill DC, Pouget A. The Bayesian brain: the role of uncertainty in neural coding and computation. Trends in Neurosciences. 2004 Dec 1;27(12):712–9.

54. Tenenbaum JB, Griffiths TL, Kemp C. Theory-based Bayesian models of inductive learning and reasoning. Trends in Cognitive Sciences. 2006 Jul 1;10(7):309–18.

55. Daunizeau J, Ouden HEM den, Pessiglione M, Kiebel SJ, Stephan KE, Friston KJ. Observing the Observer (I): Meta-Bayesian Models of Learning and Decision-Making. PLOS ONE. 2010 Dec 14;5(12):e15554.

56. Chi T, Ru P, Shamma SA. Multiresolution spectrotemporal analysis of complex sounds. J Acoust Soc Am. 2005 Aug;118(2):887–906.

57. Santoro R, Moerel M, Martino FD, Goebel R, Ugurbil K, Yacoub E, et al. Encoding of Natural Sounds at Multiple Spectral and Temporal Resolutions in the Human Auditory Cortex. PLOS Computational Biology. 2014 Jan 2;10(1):e1003412.

58. Bitterman Y, Mukamel R, Malach R, Fried I, Nelken I. Ultra-fine frequency tuning revealed in single neurons of human auditory cortex. Nature. 2008 Jan 10;451(7175):197–201.

59. Diederen KMJ, Spencer T, Vestergaard MD, Fletcher PC, Schultz W. Adaptive Prediction Error Coding in the Human Midbrain and Striatum Facilitates Behavioral Adaptation and Learning Efficiency. Neuron. 2016 Jun 1;90(5):1127–38.

60. Milne A, Zhao S, Tampakaki C, Bury G, Chait M. Sustained pupil responses are modulated by predictability of auditory sequences. J Neurosci [Internet]. 2021 Jun 3 [cited 2021 Jun 10]; Available from: https://www.jneurosci.org/content/early/2021/06/01/JNEUROSCI.2879-20.2021

61. Dalley JW, McGaughy J, O’Connell MT, Cardinal RN, Levita L, Robbins TW. Distinct Changes in Cortical Acetylcholine and Noradrenaline Efflux during Contingent and Noncontingent Performance of a Visual Attentional Task. J Neurosci. 2001 Jul 1;21(13):4908–14.

62. Yu AJ, Dayan P. Uncertainty, Neuromodulation, and Attention. Neuron. 2005 May 19;46(4):681–92.

63. Bland AR, Schaefer A. Different Varieties of Uncertainty in Human Decision-Making. Front Neurosci [Internet]. 2012 Jun 8 [cited 2018 Jun 20];6. Available from: https://www.ncbi.nlm.nih.gov/pmc/articles/PMC3370661/

64. Yarden TS, Mizrahi A, Nelken I. Context-Dependent Inhibitory Control of Stimulus-Specific Adaptation. J Neurosci. 2022 Jun 8;42(23):4629–51.

65. Natan RG, Briguglio JJ, Mwilambwe-Tshilobo L, Jones SI, Aizenberg M, Goldberg EM, et al. Complementary control of sensory adaptation by two types of cortical interneurons. eLife Sciences. 2015 Oct 13;4:e09868.

66. Natan RG, Rao W, Geffen MN. Cortical Interneurons Differentially Shape Frequency Tuning following Adaptation. Cell Rep. 2017 Oct 24;21(4):878–90.

67. Richter LMA, Gjorgjieva J. A circuit mechanism for independent modulation of excitatory and inhibitory firing rates after sensory deprivation. Proceedings of the National Academy of Sciences. 2022 Aug 9;119(32):e2116895119.

68. Schulz A, Miehl C, Berry MJ II, Gjorgjieva J. The generation of cortical novelty responses through inhibitory plasticity. Geffen MN, Gold JI, Geffen MN, editors. eLife. 2021 Oct 14;10:e65309.

69. Warren BM, Ackroff JM. Two types of auditory sequence perception. Perception & Psychophysics. 1976 Sep;20(5):387–94.

70. Warren RM. Auditory Perception: An Analysis and Synthesis [Internet]. Cambridge: Cambridge University Press; 2008 [cited 2015 Dec 24]. Available from: http://ebooks.cambridge.org/ref/id/CBO9780511754777

71. Warren RM, Obusek CJ. Identification of temporal order within auditory sequences. Perception & Psychophysics. 1972 Jan;12(1):86–90.

72. Brainard DH. The Psychophysics Toolbox. Spat Vis. 1997;10(4):433–6.

73. Kleiner M, Brainard D, Pelli D, Ingling A, Murray R, Broussard C. What’s new in psychtoolbox-3. Perception. 2007;36(14):1–16.

74. Oostenveld R, Fries P, Maris E, Schoffelen JM. FieldTrip: Open Source Software for Advanced Analysis of MEG, EEG, and Invasive Electrophysiological Data. Comput Intell Neurosci. 2010 Dec 23;2011(2011):156869.

75. de Cheveigné A, Arzounian D. Robust detrending, rereferencing, outlier detection, and inpainting for multichannel data. NeuroImage. 2018 May 15;172:903–12.

76. de Cheveigné A, Parra LC. Joint decorrelation, a versatile tool for multichannel data analysis. NeuroImage. 2014 Sep;98:487–505.

77. de Cheveigné A, Simon JZ. Denoising based on spatial filtering. Journal of Neuroscience Methods. 2008 Jun 30;171(2):331–9.

78. Efron B, Tibshirani RJ. An Introduction to the Bootstrap. New York: Chapman and Hall/CRC; 1994. 456 p.

79. Macmillan NA. Detection Theory: A User’s Guide. CUP Archive; 1991. 436 p.

80. Greenhouse SW, Geisser S. On methods in the analysis of profile data. Psychometrika. 1959 Jun;24(2):95–112.

81. Adams RP, MacKay DJC. Bayesian Online Changepoint Detection. arXiv:07103742 [stat] [Internet]. 2007 Oct 19 [cited 2019 Apr 24]; Available from: http://arxiv.org/abs/0710.3742

82. Murphy KP. Conjugate Bayesian analysis of the Gaussian distribution. 2007;1(7):29.

